# Regulation of gene expression by repression condensates during development

**DOI:** 10.1101/2020.03.03.975680

**Authors:** Nicholas Treen, Shunsuke F. Shimobayashi, Jorine Eeftens, Clifford P. Brangwynne, Michael S. Levine

## Abstract

There is emerging evidence for transcription condensates in the activation of gene expression^1–3^. However, there is considerably less information regarding transcriptional repression, despite its pervasive importance in regulating gene expression in development and disease. Here, we explore the role of liquid-liquid phase separation (LLPS) in the organization of the Groucho/TLE (Gro) family of transcriptional corepressors, which interact with a variety of sequence-specific repressors such as Hes/Hairy^4^. Gro-dependent repressors have been implicated in a variety of developmental processes, including segmentation of the *Drosophila* embryo and somitogenesis in vertebrates. These repressors bind to specific recognition sequences, but instead of interacting with coactivators (e.g., Mediator) they recruit Gro corepressors^5^. Gro contains a series of WD40 repeats that are thought to mediate oligomerization^6^. How putative Hes/Gro oligomers repress transcription has been the subject of numerous studies^5, 6^. Here we show that Hes/Gro complexes form discrete puncta within nuclei of living *Ciona* embryos. These puncta rapidly dissolve during the onset of mitosis and reappear in the ensuing cell cycle. Modified Hes/Gro complexes that are unable to bind DNA exhibit the properties of viscous liquid droplets, similar to those underlying the biogenesis of P-granules in *C. elegans*^7^ and nucleoli in *Xenopus* oocytes^8^. These observations provide vivid evidence for LLPS in the control of gene expression and suggest a simple physical exclusion mechanism for transcriptional repression. WD40 repeats have been implicated in a wide variety of cellular processes in addition to transcriptional repression^9^. We suggest that protein interactions using WD40 motifs might be a common feature of processes reliant on LLPS.

## Main Text

There is emerging evidence that gene activation is accompanied by the recruitment of large clusters of transcription complexes, particularly Mediator and RNA Polymerase II (Pol II)^1–3, 10^. However, there is controversy regarding the physical properties of these clusters^11^. Some believe that they form condensates through liquid-liquid phase separation (LLPS). But the resulting Mediator condensates are hard to visualize, highly unstable, and identified only at genetic loci regulated by super-enhancers, which represent less than 1% of all enhancers present in mammalian genomes^12^.

To explore a more general role for LLPS in gene regulation we examined the Hes/Hairy family of sequence-specific transcriptional repressors, which have been implicated in a variety of developmental processes including segmentation of the *Drosophila* embryo and somitogenesis in vertebrates^4^. These proteins recognize specific DNA sequence motifs via a basic helix-loop-helix domain, but instead of recruiting coactivators such as components of the Mediator complex, they instead interact with the Groucho/TLE (Gro) family of corepressor proteins through a short C-terminal peptide motif, WRPW^13^. Gro contains a series of WD40 repeats that have been shown to mediate the formation of Hes/Gro oligomers^5, 6^, which establish stable and dominant repression of gene activity^14^. Here, we sought to determine whether LLPS dictates this oligomerization process.

We examined this possibility using an expression assay in living *Ciona* embryos, taking advantage of the large nuclei (8-10 microns in diameter) and ease of expressing fluorescent fusion proteins by simple electroporation assays^15^ (summarized in Fig. 1A). The *Sox1/2/3* enhancer mediates expression in ectodermal cells^16^ by the onset of gastrulation at the 110-cell stage (Fig. 1B). These cells are particularly suitable for analysis of intracellular dynamics as they do not undergo the complex movements seen for the presumptive endoderm and mesoderm located on the other (vegetal) side of gastrulating embryos^17^. Coding sequences of interest were placed downstream of the *Sox1/2/3* enhancer, and fluorescent moieties such as mNeongreen^18^ (mNg, 236 amino acid residues) were fused in-frame in either the 5’ or 3’ position. There is only a ∼3-fold increase in the levels of expression as compared with endogenous *Sox 1/2/3* products (Extended Data Fig. 1). As a proof of principle, we examined the distribution of the Fibrillarin (Fbl) protein, an integral component of the nucleolus^19^. As shown previously for nucleoli in *Xenopus* oocytes^8^, *Ciona* nucleoli display properties of viscous liquid droplets that undergo variable fusions (Extended Data Fig. 2, Supplementary Videos 1, 2).

**Fig. 1:**
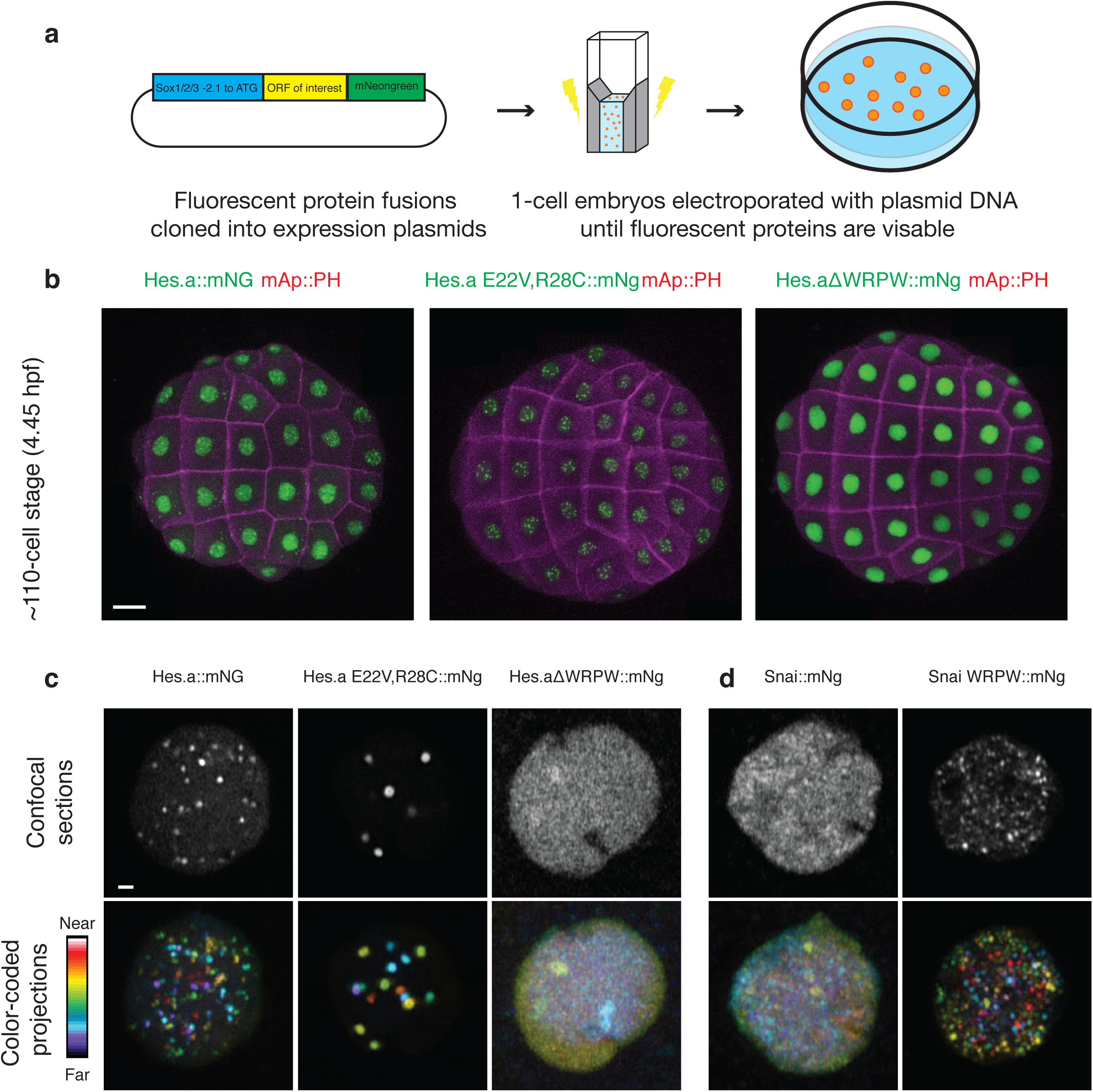
The Hes.a repressor forms puncta in *Ciona* embryos dependent on DNA binding and the presence of a WRPW domain. **a**, Schematic of the electroporation procedure used to transfect *Ciona* embryos with plasmid DNA. **b**, Maximum intensity confocal projections of ∼110-cell stage embryos expressing transgenes from pSP *Sox1/2/3* plasmids. Cell membranes are colored magenta and Hes.a::mNg fusion proteins are green. The embryos are oriented to show the animal hemisphere, anterior left. Scale bar = 20 μm. **c**, Confocal images of individual nuclei expressing Hes.a proteins fused to mNg. Single confocal sections are shown in white, color coded projections are shown with the indicated look up table **d**, Same as **c** but for the Snai::mNg. Scale bar = 1 μm.

**Fig. 2:**
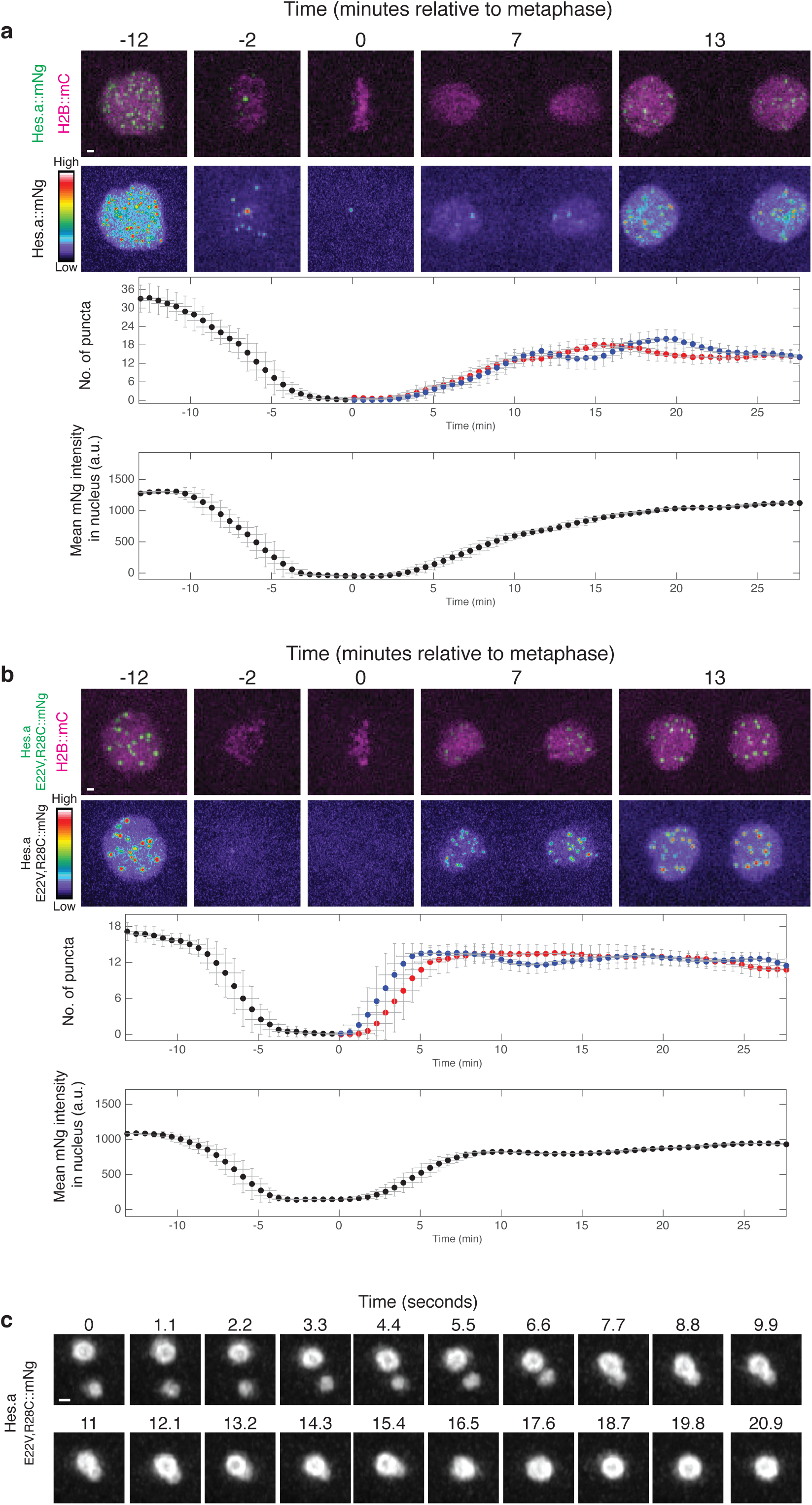
Hes.a shows liquid-like properties throughout the cell cycle. a, Time-lapse maximum intensity projection confocal images of a single *Ciona* nucleus from the 7th to 8th mitosis. Hes.a::mNg is shown in green and with the indicated look up table. H2B::mCh is shown in magenta. Graphs are depicting properties of green fluorescence within the red fluorescence region. Error bars show the standard deviation ± 100 sec. Scale bar = 1 μm. **b**, Same as A but for the Hes.a E22V,R28C mutant. **c,** Time-lapse maximum intensity projection confocal images of the fusion of 2 Hes.a E22V,R28C::mNg puncta. Scale bar = 0.5 μm.

Using this assay, we found that *Ciona* Hes.a protein is distributed in multiple puncta per nucleus (Fig. 1B). It is likely that these puncta are formed by Hes.a-Gro interactions at localized sites within the *Ciona* genome containing clusters of Hes.a binding sites. We investigated the properties of these puncta to see how closely they resemble liquid-liquid phase separated condensates, such as nucleoli. Particular efforts focused on two different Hes.a protein variants (Fig. 1B,C). The first contains two amino acid substitutions (E22V and R28C) in the bHLH domain that eliminate DNA binding^20^ while the other lacks the WRPW peptide motif at the C-terminus that is essential for interactions with Gro^5^. The loss of DNA binding leads to the formation of large Hes.a puncta, whereas loss of interactions with Gro causes the opposite phenotype—virtual elimination of puncta (Fig. 1B, C). Remarkably, the WRPW peptide motif is sufficient to confer clustering of DNA binding proteins that normally display dispersed distribution profiles, such as Snail where the addition of the WRPW motif induces the formation of Snail puncta (Fig. 1D, Extended Data Fig. 3).

**Fig. 3:**
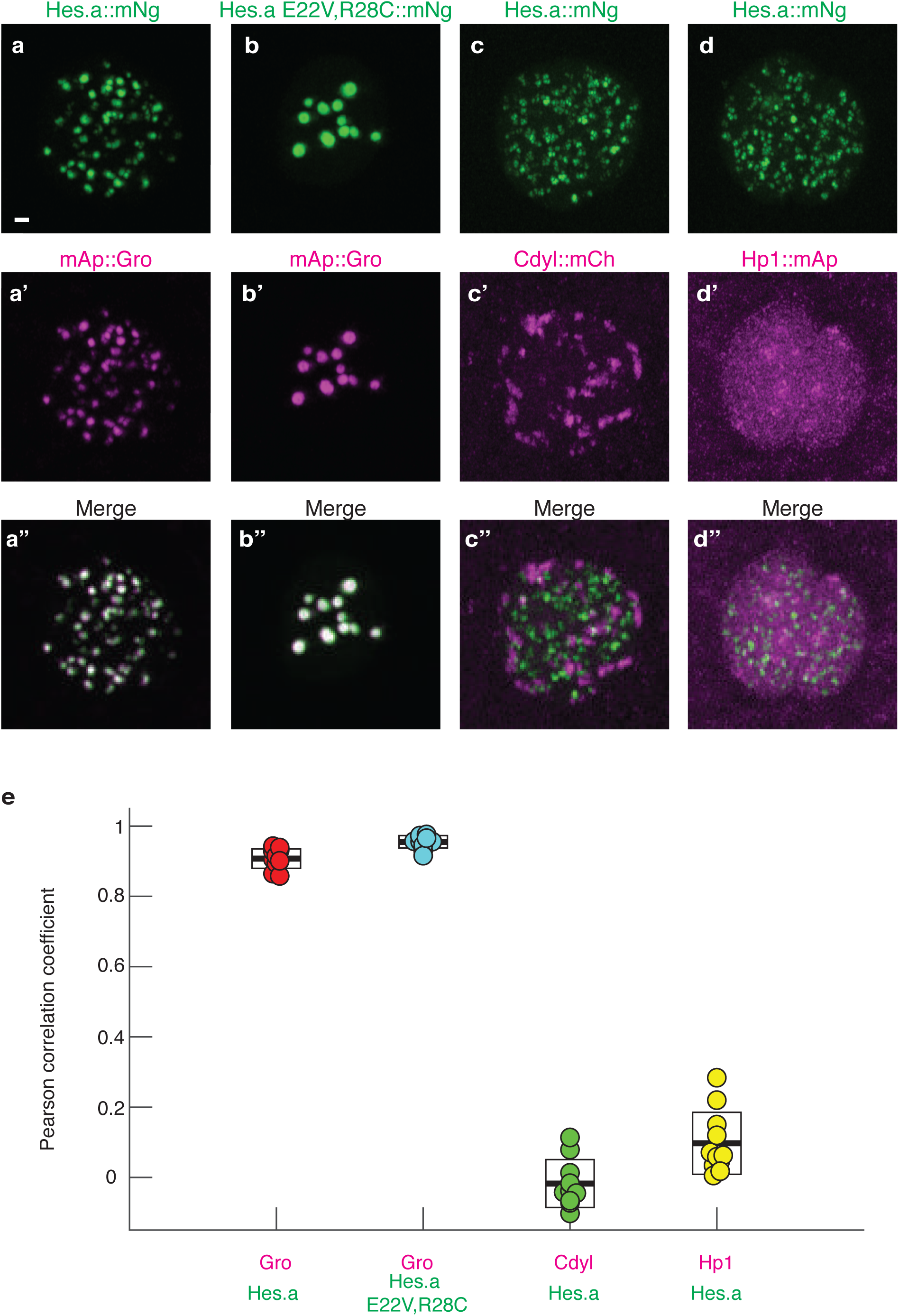
Hes.a/Groucho puncta are a novel molecular condensates. a, Maximum intensity projection confocal images of single *Ciona* nuclei expressing Hes.a::mNg. a’, mAp::Gro Fluorescence. a’’, The merged green and red channels for a and a’. Scale bar = 1 μm. b-b’’, As for the a series but for Hes.a E22V,R28C::mNG and mAp::Gro Green and red channels are shown individually and merged. c-c’’, Hes.a::mNg and Cdyl:mCh d-d’’, Hes.a::mNg and Hp1::mAp c, Pearson correlation coefficients of the experiments shown in a-d’’. Boxes display the mean and standard deviations.

To determine whether Hes.a/Gro puncta correlate with transcriptional repression we examined the activities of a *ZicL>H2b::mCherry* (mCh) reporter gene (Extended Data Fig. 4). ZicL is an authentic target of the Hes.a repressor in early development^21^. Wild-type Hes.a efficiently represses the ZicL reporter, whereas mutant forms that are unable to bind DNA or interact with Gro do not (Extended Data Fig. 4). These results suggest that the formation of the Hes.a puncta, binding to both DNA and Gro, are required for repression.

**Fig. 4:**
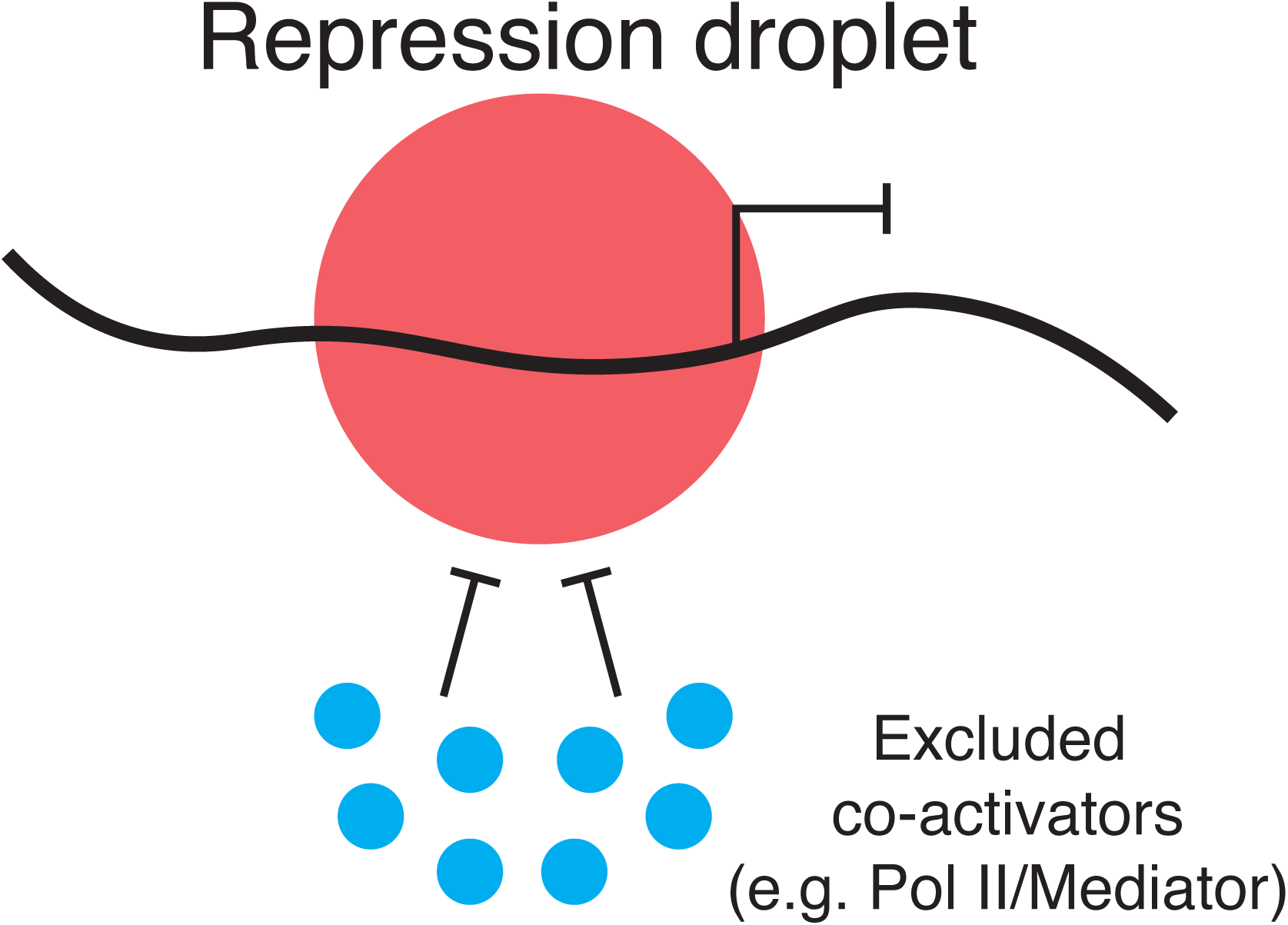
Transcriptional repression through stable sequestration of regulatory DNAs. This schematic depicts an example where transcriptional activation in inhibited by the formation of a liquid repression droplet (red) upon a regulatory region of DNA. Transcriptional activators (blue) are excluded from this droplet.

We next examined the dynamics of Hes.a puncta to determine if they display liquid-like properties associated with LLPS. Both the wild-type and DNA binding mutant (E22V,R28C) produce puncta that are detected throughout interphase, but are abolished during mitosis before reforming in daughter nuclei (Fig. 2A, B Supplementary Videos 3, 4). The mutant exhibits more rapid dynamics than the wild-type protein, it dissolves more quickly during mitosis and re-forms more rapidly in daughter nuclei following mitosis (Fig. 2B, Supplementary Videos 3, 4). These results raise the possibility that the binding of Hes.a to its cognate DNA recognition sequences could localize phase separation to specific nanoscopic regions of the genome. When DNA binding is disrupted, the resulting Hes.a/Gro puncta display liquid properties that are difficult to observe for wild-type puncta.

There is some controversy concerning the criteria underlying the formation of biomolecular condensates via LLPS^11, 22, 23^. However, one critical property is dynamic fusions of individual droplets^7, 8^. Such fusions are readily detected for the E22V,R28C mutant (Fig. 2C, Supplementary Video 5), but not for the wild-type protein. However, there is a progressive reduction in the number of wild-type Hes.a/Gro puncta during multiple cell cycles without a corresponding diminishment in fluorescence intensity (Fig. 2A). A possible explanation for this observation is that wild-type puncta undergo fusion events as nuclei diminish in size, creating higher concentrations of compact chromatin as compared with earlier stages of development.

Previous studies have shown that heterochromatin is compartmentalized within the nucleus. HP1 binds constitutive heterochromatin (H3K9me3)^24, 25^ and coalesces in living *Drosophila* embryos and cells to form several large condensates located near the periphery of the nucleus^26, 27^. Polycomb repression complexes (e.g., PRC2) bind to facultative heterochromatin (H3K27me3)^28^ and also form higher order puncta resembling condensates^29^. Double labeling assays were used to determine whether Hes.a/Gro condensates are associated with either type of heterochromatin (Fig. 3). Control experiments showed the co-localization of Hes.a and Gro fusion proteins (Fig. 3 A,B,E). Co-expression of the wild-type Hes.a::Ng fusion protein with Fbl (nucleoli), Cdyl (a protein directly associated with both PRC2 and H3K27me3), or HP1 fusion proteins reveals little or no significant co-localization of Hes.a and heterochromatin or nucleoli (Fig. 3 C,D,E; Extended Data Fig. 5). These observations suggest that Hes.a does not silence gene expression by associating with heterochromatin, although it shares the property of forming condensates.

We have presented evidence that Hes.a/Gro complexes form condensates through LLPS. These condensates are likely to depend on dynamic Hes.a-Gro interactions since neither protein alone forms puncta. Gro proteins contain oligomerization and disordered domains in addition to the WD40 repeats (Extended Data Fig. 6). These domains have been implicated in the formation of extended oligomers along the chromatin template^6^. There is emerging evidence for the role of coupling oligomerization with protein disorder to drive phase separation^30^. We suggest that interactions between the oligomerization and disordered domains of Hes.a and Gro induce LLPS to trigger the formation of condensates, similar to other protein and nucleic acid-rich condensates such as P granules and nucleoli^31, 32^. However, in this context the growth and coarsening of these condensates appears to be limited by DNA binding since the E22V,R28C Hes.a mutant displays conspicuous fusion events producing considerably larger condensates as compared with wild-type complexes.

Hes.a/Gro condensates are considerably more stable than putative activation condensates, which typically display short half-lives of just ∼10 seconds, although a small subset persist for minutes^1^. In contrast, Hes.a/Gro condensates are longer lived, and more evocative of nucleoli.

Once formed, they persist throughout the cell cycle and do not dissolve until mitosis. We do not detect stable condensates for a variety of sequence-specific activators that were tested in our *Ciona* embryo assay (Extended Data Fig. 3). However, some can form puncta upon addition of a WRPW motif that mediates interactions with Gro (Extended Data Fig. 3). Human Hes/TLE complexes also form condensates in cultured cells. To test the concept that oligomerization can drive phase separation of these proteins we utilized the recently developed corelet optogenetic system^30^. In cultured human cells, Hes1 and TLE corelets formed colocalized puncta upon light activation (Extended Data Fig. 7A,B; Supplementary Videos 6, 7). A Hes1 DNA binding mutant (E43V,R49C) produces droplets that are more dynamic than the normal Hes1 protein, similar to the behavior of the *Ciona* Hes.a E22V,R28C mutant (Extended Data Fig. 7C, Supplementary Video S8). Since previous work has shown that corelet-induced puncta exhibit hallmarks of phase separation, these data provide further support for Hes.a driving repressive condensates through LLPS.

Gro has 7 WD40 motifs that are required for the formation of repressive condensates^5^. These motifs are a common feature of multi-protein complexes that are known or suspected to undergo LLPS^33, 34^. WD40 proteins are involved in a variety of cellular processes such as cell signaling and DNA repair, in addition to transcriptional repression as described above^9^. We propose that interactions between proteins containing disordered domains with those containing WD40 repeats might be a key trigger for the oligomerization of biological condensates. In fact, it seems likely that Polycomb repression bodies (see Fig. 3) may be formed by LLPS since the EED subunit of the PRC2 complex contains 7 WD40 repeats, as seen for Gro^35^.

We propose that repression condensates inhibit gene expression by the mechanical exclusion of transcriptional activators, coactivator complexes such as Mediator, or active chromatin^36^ (Fig. 4). These repression condensates might correspond to the inactive, B compartments observed in Hi-C contact maps^37^. It is interesting that HP1 and Polycomb also form stable condensates^26, 27, 29^. The long-term stability of these condensates within a cell cycle is consistent with the dominance of transcriptional repression in the control of gene expression^14^. The dissolution of repressive condensates at mitosis may be a pre-requisite for activating new programs of gene expression during development.

## Acknowledgements

We thank members of the Levine and Brangwynne labs for their support, especially Laurence Lemaire for sharing reagents, Chen Cao for providing raw single cell gene expression data, and Evangelos Gatzogiannis for help with imaging. This research was funded by NIH grants (NS076542 to MSL; 01 DA040601 to CPB) and the HHMI (to CPB). NT is funded by a Princeton Catalysis Initiative grant (to MSL and CPB). JE is funded by an NWO Rubicon grant.

## Author Contributions

NT and MSL conceived the project and designed the experiments. NT performed the *Ciona* experiments. SFS performed image analysis. JE designed and performed human cell experiments. NT and MSL wrote the paper with input from all other authors.

## Supplementary Information

### Methods

#### Animals

Wild type *Ciona intestinalis* (Type A, also recently referred to as *Ciona robusta*) sourced from San Diego County, Ca were supplied by M-Rep. Animals were kept in aerated artificial seawater at 18°C. All procedures involving live animals were performed at ∼18°C.

#### Human Cells

Corelet containing cells were all HEK293. Transfected by lentivirus as previously described^30^.

#### Molecular cloning

The upstream regulatory region of Sox1/2/3 from the translation initiation site to 2.3 kb upstream has previously been described to activate transgene expression in the ectoderm^16^. This sequence was subcloned into pSP plasmids and the open reading frame of the gene of interest was amplified by PCR (see Supplementary Table 1) using a proofreading polymerase (Primestar, Takara). The open reading frames were fused to in frame to fluorescent protein coding sequences separated by the linker sequence: GGSGGGSGG. Plasmids were assembled from linear PCR products by treatment with NEBuilder HiFi DNA Assembly Master Mix (New England Biolabs). Full plasmid sequences and descriptions of the individual cloning steps can be provided upon request. The ZicL>H2B::mCherry plasmid has previously been described^38^ Corelet plasmids were constructed by ligating human Hes1 and TLE cDNAs into previously described plasmids for lentiviral transfection^30^.

#### Electroporation

Dechorionated Ciona Zygotes were electroporated at 30 minutes post fertilization using standard electroporation settings^15^. 30 ug of plasmid DNA was electroporated for each individual plasmid used except for *ZicL>H2B::mCherry* where 20 ug was electroporated.

#### Gene expression levels

Single cell gene expression levels were taken from a recently published dataset^39^. qPCR assays to determine relative levels of gene expression were performed as previously described^40^ with the exception that cDNA synthesis was performed using an iScript gDNA Clear cDNA Synthesis Kit (Biorad) with a DNase digestion following the manufacturer’s instructions. Control reactions performed without reverse transcriptase showed several hundred-fold reductions in amplification suggesting minimal contamination from genomic or plasmid DNA. Primers used are listed in Supplementary Table 1.

#### Imaging

*Ciona* embryos were imaged using a Zeiss LSM 880 inverted confocal microscope (Carl Zeiss). Embryos were mounted on 3.5cm glass bottom dishes (MatTek cat # P35G-1.5-20-C) as previously described^41^ Whole embryos were imaged using a 40x 1.2 NA C-apochromat water immersion objective. Other images were taken with a 63x 1.4 NA plan-apochromat oil immersion objective. All imaging was performed using an Airyscan detector in fast mode.

Images were processed using ZEN software (ZEN Version 2.3 and 2.6, Zeiss). Human cell imaging was performed using a Nikon A1 laser scanning confocal microscope equipped with a CO_2_ microscope stage incubator under 5% CO_2_ and 37°C with a plan-apochromat 60X 1.4 NA oil immersion objective.

#### Image analysis

The colocalization between green and red fluorescent channels was quantified with pixel-based intensity correlation pearson correlation coefficients^42^ where 1 is a perfect correlation, 0 is no correlation and -1 is perfect anti-correlation.

**Extended Data Fig. 1:**
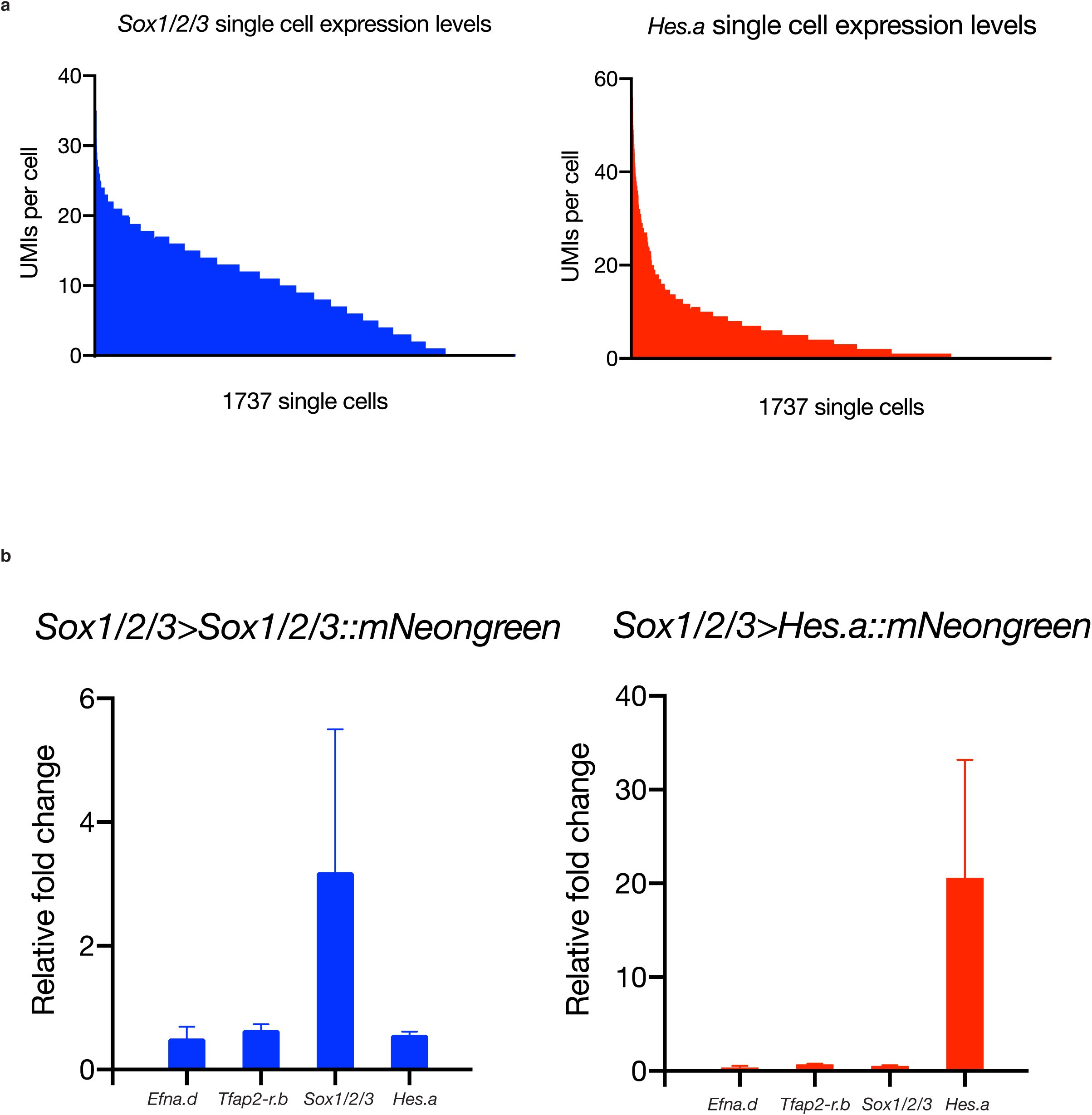
*Sox1/2/3* and *Hes.a* overexpression levels. **a**, Single cell expression levels for *Sox1/2/3* and *Hes.a* at the 110-cell stage from a previously published dataset^39^. Each UMI represents an individual transcript and can be used as an approximation for number mRNAs per cell. **b**, Relative changes in mRNA levels of 4 transcription factors from hundreds of pooled embryos at the 110-cell stage after electroporation with *Sox1/2/3>Sox1/2/3::mNg* or *Sox1/2/3>Hes.a::mNg*. Bars depict mean and standard errors.

**Extended Data Fig. 2:**
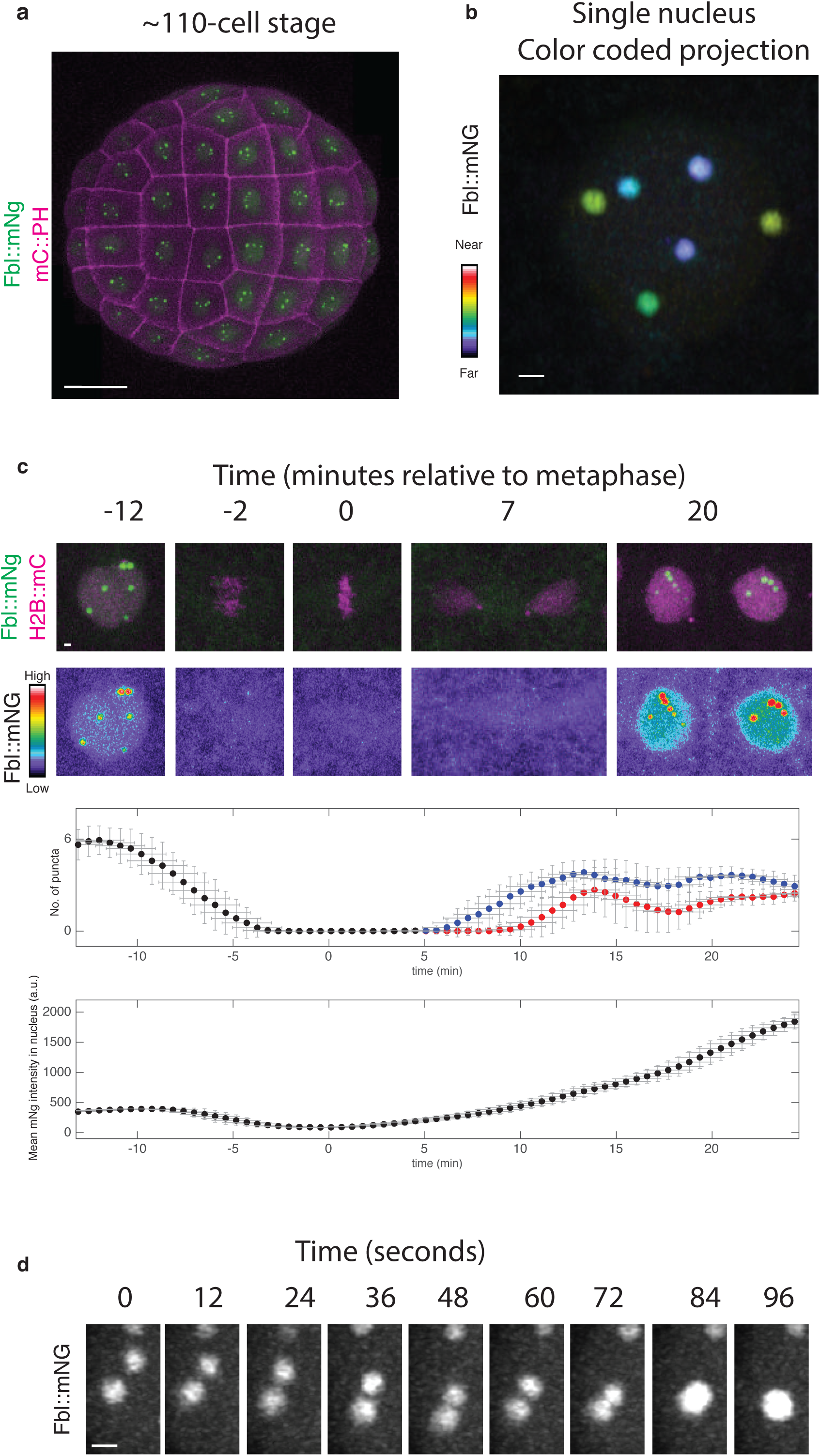
The nucleoli of *Ciona* embryos show liquid properties. **a**, Confocal maximum intensity projection of a whole *Ciona* embryo expressing a Fbl::mNg transgene Scale bar = 20 μm. **b,** Color coded projections of a single nucleus expressing Fbl::mNg. Scale bar = 1 μm. **c,** Time-lapse maximum intensity projection confocal images of a single *Ciona* nucleus from the 7th to 8th mitosis. Fbl::mNg is shown in green and with the indicated look up table. H2B::mCh is shown in magenta. Graphs depict properties of green fluorescence within the red fluorescence region. Error bars show the standard deviation ± 100 sec. Scale bar = 1 μm. **d**, Time-lapse maximum intensity projection confocal images of the fusion of 2 nucleoli. Scale bar = 0.5 μm.

**Extended Data Fig. 3:**
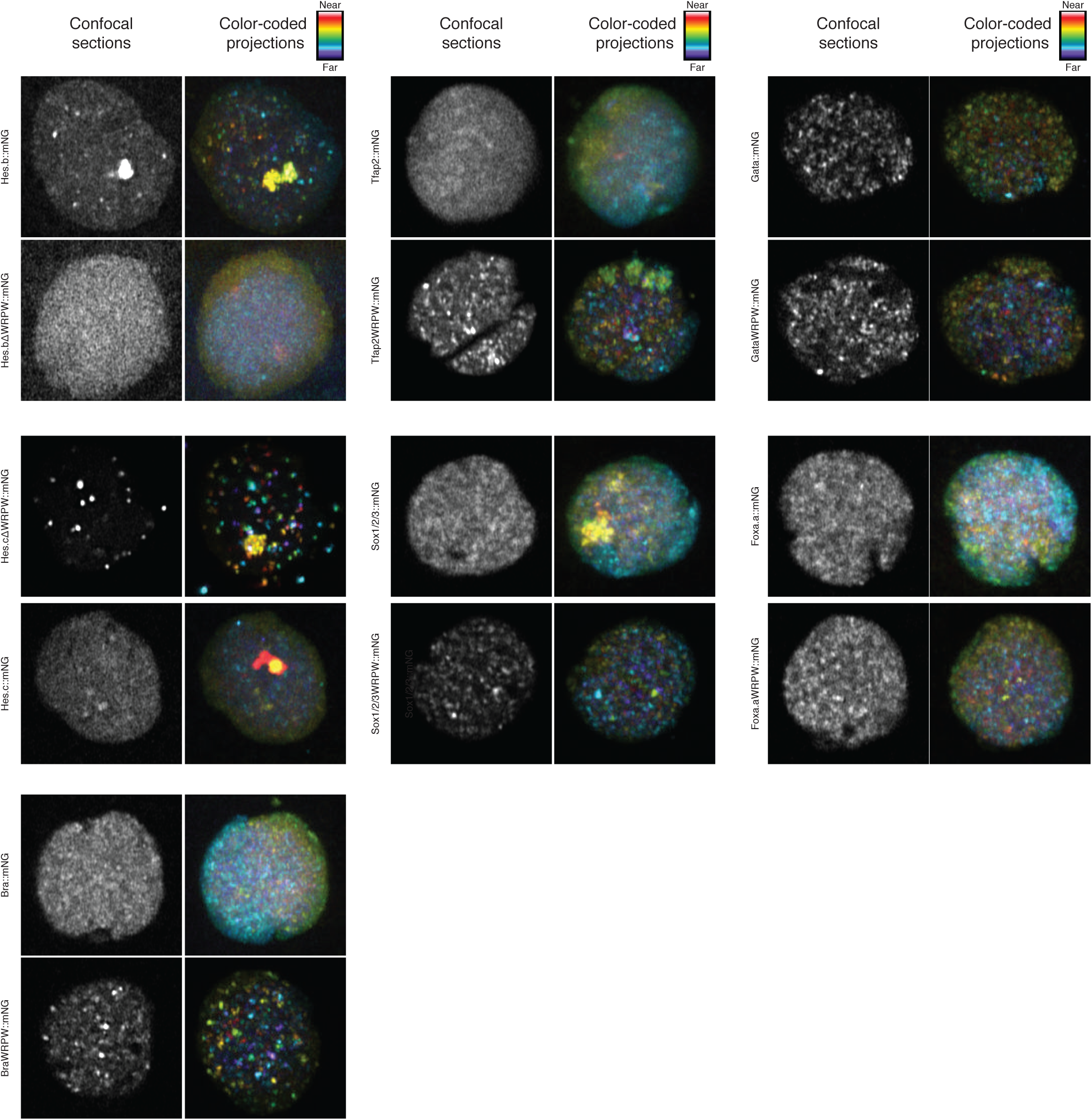
The WRPW motif is required for and can induce transcription factors to from puncta. Single confocal sections and color-coded projections of single nuclei expressing transcription factors fused to mNg with or without a C-terminal WRPW domain. Scale bar = 1 μm.

**Extended Data Fig. 4:**
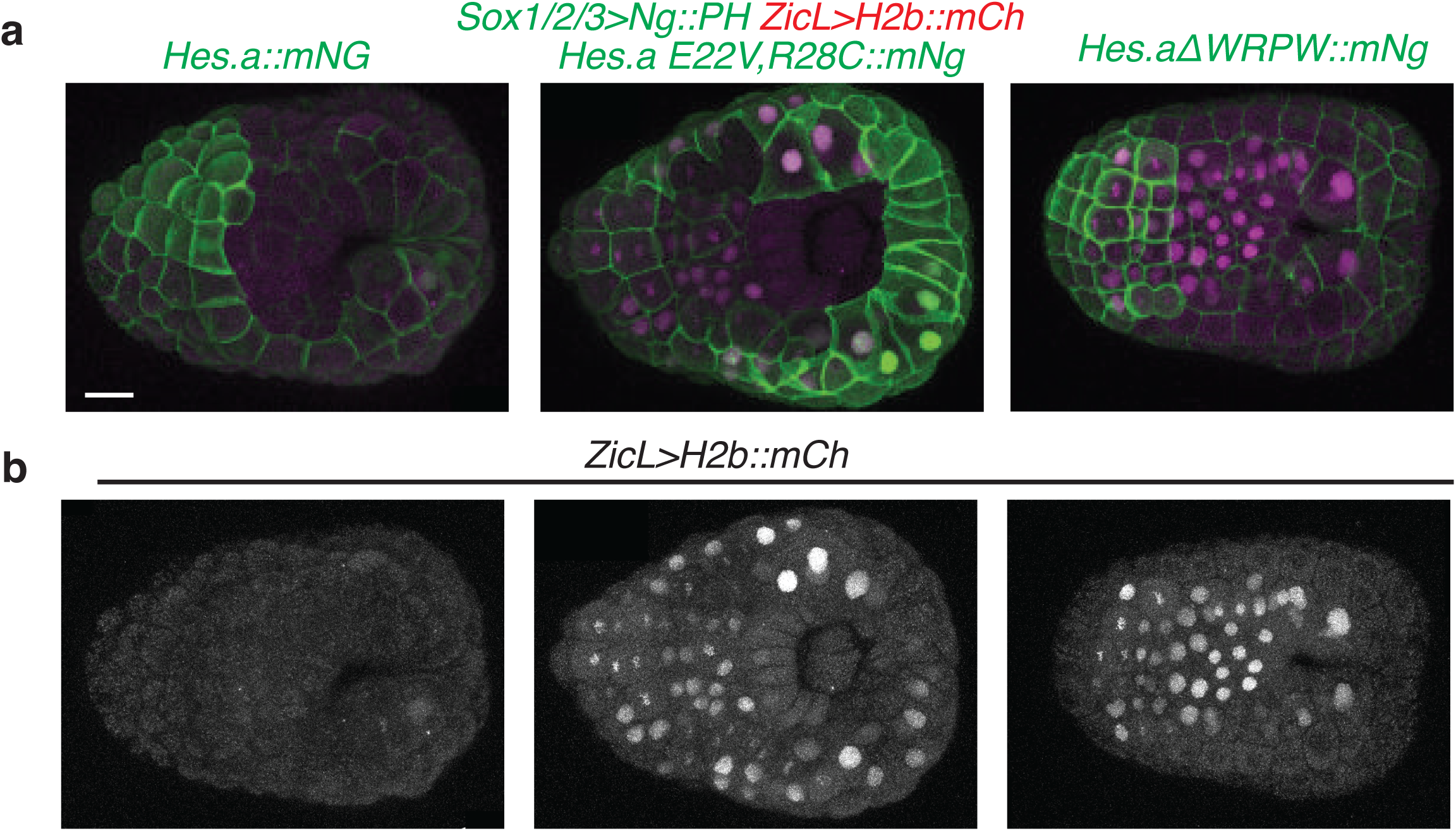
Hes.a represses *ZicL*. **a**, Maximum intensity confocal projections of gastrula stage *Ciona* embryos. Hes.a and mNG::PH fusion proteins are in green expressed by the *Sox1/2/3* regulatory region. H2B::mCherry is in magenta expressed by the *ZicL* regulatory region. Scale bar = 20 μm **b**, The red fluorescence channel from **a** for *ZicL>H2B::mCh* shown in grayscale.

**Extended Data Fig. 5:**
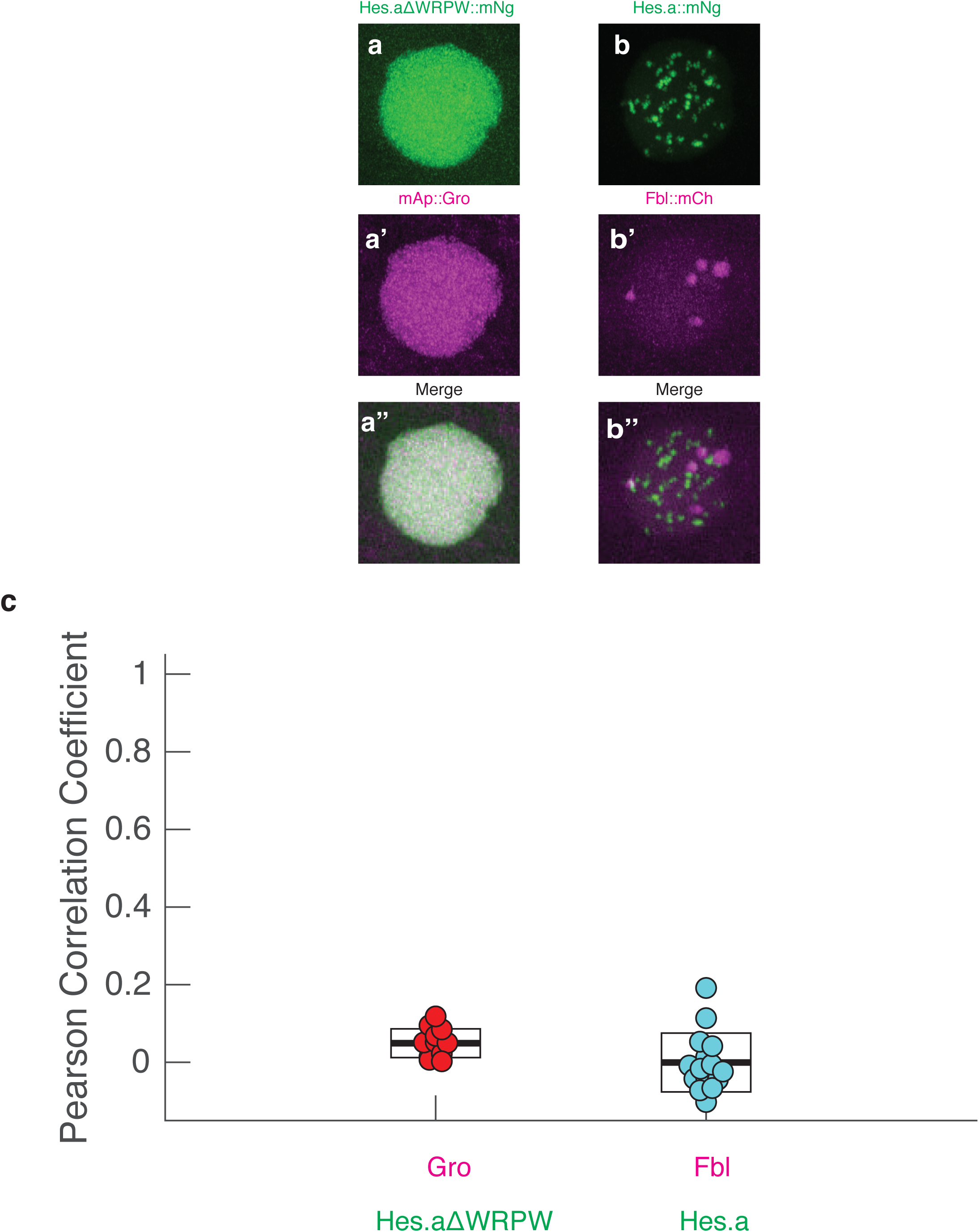
Hes.a colocalization analysis. **a**, Maximum intensity confocal projections of *Ciona* nuclei expressing Hes.a aΔWRPW::mNG. Scale bar = 1 μm. **a**’ As **a** but for mAp::Gro. **a**’’, Merged green and red channels for **a** and **a**’. **b-b**’’, Same as **a**-**a**’’ but for Hes.a::mNg and Fbl::mCh. **C**, Person correlation coefficients of the experiments shown in **a**-**b**’’. Boxes display the mean and standard deviations.

**Extended Data Fig. 6:**
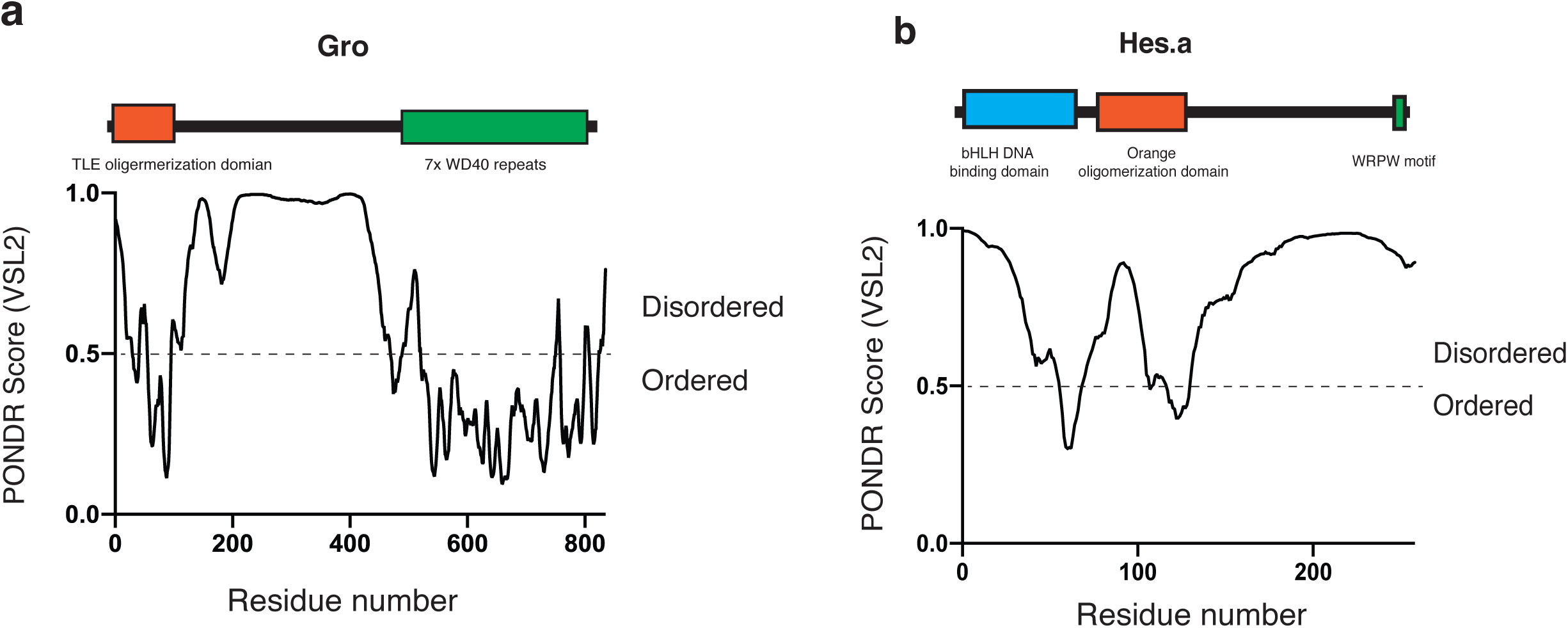
Primary structure of Groucho and Hes.a proteins. **a,** For Gro, annotated domains are indicated as well as a PONDR score to show predictions of ordered or disordered structure throughout the protein. **b,** Same analysis as **a** but for Hes.a.

**Extended Data Fig. 7:**
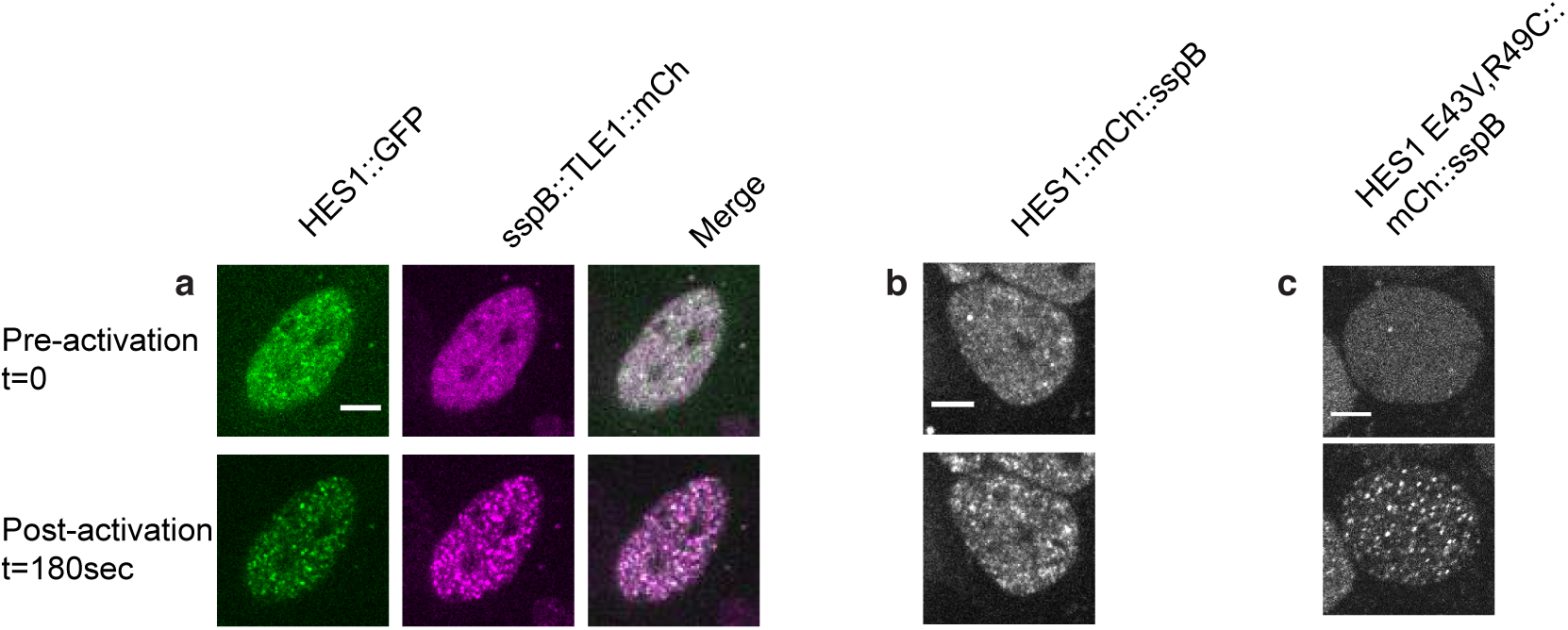
Human Hes1 and TLE corelets show light inducible aggregation and phase separation. **a**, a single human cell nucleus expressing Hes1::GFP and sspB::mCh-TLE1 before and after light activation. **b**, Wildtype Hes1-mCh-sspB behavior before and after light induced binding of corelets. **c**, Same as **b,** but for the DNA binding domain mutant of Hes1. Scale bar = 5 μm

**Supplementary Table 1:**
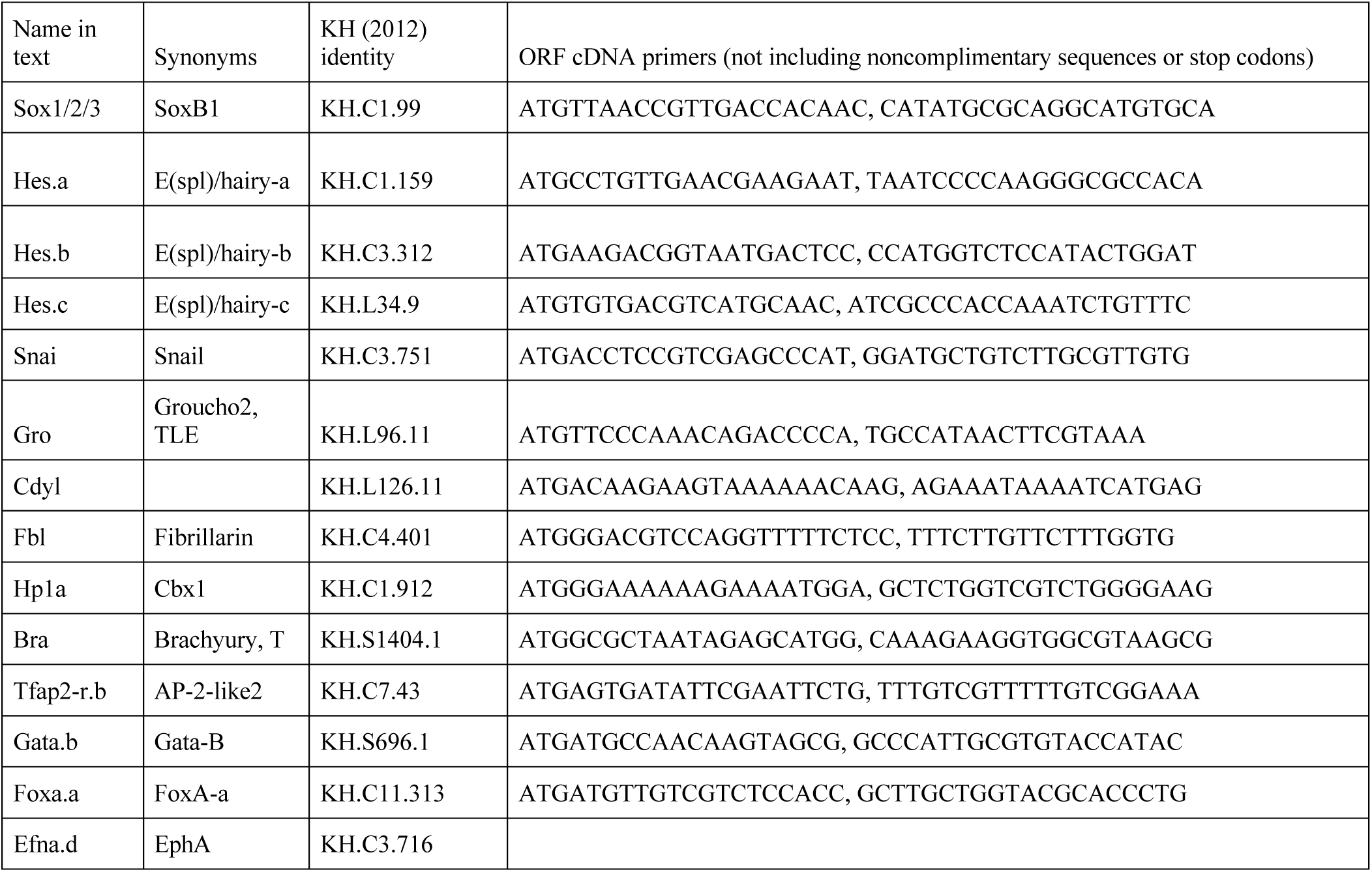

**Video 1: Fibrillarin dynamics during development.**

Maximum intensity confocal projection of Fbl::mNg (in green) and H2b::mCh (in magenta) throughout a full mitosis. Time is in minutes relative to metaphase.

**Video 2: The fusion of 2 fibrillarin droplets.**

Maximum intensity confocal projection of the fusion of 2 Fbl::mNg droplets.

**Video 3: Hes.a dynamics during development.**

Maximum intensity confocal projection of Hes.a::mNg (in green) and H2b::mCh (in magenta) throughout a full mitosis. Time is in minutes relative to metaphase.

**Video 4: Hes.a DNA binding mutant dynamics during development**

Maximum intensity confocal projection of Hes.a E22v,R28C::mNg (in green) and H2b::mCh (in magenta) throughout a full mitosis. Time is in minutes relative to metaphase.

**Video 5: The fusion of 2 Hes.a E22V,R28C droplets.**

Maximum intensity confocal projection of the fusion of 2 Hes.a E22v,R28C::mNg droplets.

**Video 6: Hes1 TLE corelet colocalization.**

A single human cell nucleus expressing Hes1::GFP (in green) and sspB::mCh-TLE (in magenta) Time is in seconds after activation. Scale bar = 5 μm

**Video 7: Hes1 corelet aggregation.**

A single human cell nucleus expressing wildtype Hes1-mCh-sspB Time is in seconds after activation. Scale bar = 5 μm.

**Video 8: Hes1 E43V,R49C mutant corelet aggregation.**

A single human cell nucleus expressing Hes1E43V,R49C-mCh-sspB Time is in seconds after activation. Scale bar = 5 μm.

